# In vivo complex haploinsufficiency-based genetic analysis identifies a transcription factor circuit regulating *Candida albicans* oropharyngeal infection and epithelial cell endocytosis

**DOI:** 10.1101/2021.05.24.445409

**Authors:** Norma V. Solis, Rohan S. Wakade, Tomye L. Ollinger, Melanie Wellington, Aaron P. Mitchell, Scott G. Filler, Damian J. Krysan

**Author notes:** Corresponding Author: Damian J. Krysan, 2040 Med Labs 25 S. Grand Avenue, Department of Pediatrics and Microbiology/Immunology, Carver College of Medicine, University of Iowa, Iowa City Iowa 52242.

## Abstract

Oropharyngeal candidiasis (OPC) is a common infection that complicates a wide range of medical conditions which can cause either mild or severe disease depending on the patient. The pathobiology of OPC shares many features with candidal biofilms of abiotic surfaces. The transcriptional regulation of *C. albicans* formation of biofilms on abiotic surfaces has been extensively characterized and involves six key transcription factors (Efg1, Ndt80, Rob1, Bcr1, Brg1, and Tec1). To determine whether this same in vitro biofilm transcriptional regulatory network played a role in OPC, we have carried out the first systematic genetic interaction analysis in a mouse model of *C. albicans* infection. Whereas all six transcription factors are required for in vitro biofilm formation, only three homozygous deletion mutants (*tec1*ΔΔ, *bcr1*ΔΔ, and *rob1*ΔΔ) and one heterozygous mutant (*tec1*Δ/*TEC1*) have reduced infectivity in a mouse model of OPC, indicating the network is more robust in vivo than in vitro. Although single mutants (heterozygous or homozygous) of *BRG1* and *EFG1* have no effect on fungal burden, the double heterozygous and homozygous mutants have dramatically reduced infectivity, indicating a critical genetic interaction between these two transcription factors. Using epistasis analysis, we have formulated a genetic circuit [*EFG1*+*BRG1*]→*TEC1*→*BCR1* that is required for OPC infectivity and oral epithelial cell endocytosis. Surprisingly, we also found transcription factor mutants with in vitro defects in filamentation such as *efg1*ΔΔ and *brg1*ΔΔ filament during oral infection and that decreased filamentation did not correlate with decreased infectivity. Taken together, these data indicate that key in vitro biofilm transcription factors are involved in OPC but that the network characteristics and functional connections are remodeled significantly during interactions with tissues.

**Author Summary:** The pathology of oral candidiasis has features of biofilm formation, a well-studied process in vitro. Based on that analogy, we hypothesized that network of transcription factors that regulates in vitro biofilm formation might have similarities and differences in during oral infection. To test this, we employed the first systematic genetic interaction analysis of *C. albicans* in a mouse model of oropharyngeal infection. This revealed that the six regulators involved in in vitro biofilm formation played roles in vivo but that the functional connections between factors were quite distinct. Surprisingly, we also found that, while many of the factors are required for filamentation in vitro, none of the transcription factor deletion mutants was deficient for this key virulence trait in vivo. These observations clearly demonstrate that *C. albicans* regulates key aspects of its biology differently in vitro and in vivo.

## Introduction

*Candida albicans* is a commensal component of the human mycobiome that frequently causes mucosal infections of both immunocompetent and immunocompromised people [1]. The oral cavity is an important *C. albicans* niche and, consequently, this organism causes both mucosal and dental infections under specific conditions [2]. The infections of the oral mucosa caused by *C. albicans* range from minor, superficial infections such as neonatal thrush to extensive pharyngeal and, after extension, esophageal infections. Oropharyngeal candidiasis (OPC) complicates a variety of medical conditions including diabetes, radiation therapy for head and neck cancers, and alterations in saliva production [3]. The most severe infections frequently occur in patients who have altered T cell immunity. For example, people living with HIV still frequently developed OPC in the era of highly effective retroviral therapy [4]. In addition, specific primary immune deficiencies such as chronic mucocutaneous candidiasis and others lead to persistent OPC [5]. Furthermore, the development of immunomodulating therapies in other areas of medicine continues at a brisk pace, leading to increasing numbers of patients who are at-risk for fungal diseases such as OPC. Therefore, understanding the pathobiology of this relatively common fungal infection is not only of fundamental importance to fungal pathogenesis but also could inform the development of innovative approaches to its treatment.

One of the cardinal clinical features of OPC is the development of tissue adherent plaques of yeast and hyphae-stage fungi [2,3]. Histological analysis of these lesions is consistent with *C. albicans* adopting features of biofilm-phase growth such as surface adherent replication and extracellular matrix formation [6]. Results from a number of investigations including from our groups have supported this general model at both the macro- and molecular level. For example, the transcriptional regulator of in vitro biofilm formation, Bcr1 [7], is also required for infection and virulence in a mouse model of OPC [8]. In recent years, large-scale, systematic genetic studies have identified a set of *C. albicans* transcription factors (TF) that are required for in vitro biofilm formation [9, 10]. Recently, we have also used a complex haploinsufficiency-based approach to identify key interactions between the components of this transcriptional network during different stages of in vitro biofilm formation [11, 12].

To further explore the overlap between the regulation of in vitro biofilm formation and oral infection, we asked which members of the in vitro biofilm transcriptional regulatory network are required to establish oral infections [9]. In addition, we used our array of 12 double heterozygous transcription factor (TF) deletion strains to identify pairs that were critical for OPC [11]. Interestingly, although all six of the TFs we tested were absolutely required for in vitro biofilm formation, only strains containing homozygous deletions of *TEC1* or *BCR1* reduced *C. albicans* fungal burdens in our model of OPC. Genetic interaction analysis revealed that, although deletion of neither *BRG1* nor *EFG1* led to a significant reduction in infectivity, the double heterozygous strain (*efg1*Δ/*EFG1 brg1*Δ/*BRG1*) was highly attenuated; in addition, double homozygous *efg1*ΔΔ *brg1*ΔΔ strains were completely cleared from the oral cavity. This strong genetic interaction was specific to OPC because the virulence of the *efg1*Δ *brg1*Δ double heterozygous mutant was indistinguishable from WT in a model of disseminated candidiasis. Our genetic data indicate that Efg1 and Brg1 are critical for activation of the Tec1-Bcr1 axis and for the ability of *C. albicans* to invade oral epithelial cells. Thus, we have used the first in vivo interaction analysis of a TF network to identify a critical regulatory circuit for *C. albicans* oral infection.

## Materials and Methods

### Results

#### The transcriptional network for establishment of OPC is distinct from in vitro biofilm formation

In 2012, Nobile et al. characterized a network of six TFs [9] that are essential for the formation of biofilms in vitro (Fig. 1A). As part of these studies, they also showed that the TFs were, by and large, required for in vivo biofilm formation using rat models of vascular catheter and denture infections; *bcr1*ΔΔ showed a less pronounced effect in the denture model than in the catheter model. Both of the in vivo models studied by Nobile et al. involve the formation of a *C. albicans* biofilm on an inanimate surface within the host [9]. As discussed above, *C. albicans* shows features of biofilm stage growth in OPC [6]. As such, an OPC-related biofilm represents the adhesion to, and infection of, a biological, mucosal surface rather than an abiotic surface. To determine the role of in vitro biofilm TFs in a murine model of OPC, we infected corticosteroid-treated outbred CD-1 mice with both homozygous and heterozygous deletion mutants of the six biofilm TFs using the well-established model developed previously. The tissue fungal burdens were determined 5 days after inoculation. As shown in Fig. 1B, three homozygous deletion mutants, *tec1*ΔΔ, *rob1*ΔΔ and *bcr1*ΔΔ, showed a reduction in tissue fungal burden; the *tec1*Δ/*TEC1* heterozygous mutant was the only haploinsufficient strain (Fig. 1C). Bcr1 has been previously shown to be required for OPC but the roles of Tec1 and Rob1 have not been reported [8]. Thus, a set of three in vitro biofilm-related transcription factors are required for OPC infectivity in a mouse model whereas three others (Ndt80, Efg1, and Brg1) do not alter OPC infectivity. Importantly, Nobile et al. found that all six regulators were required to form normal biofilms in rodent models of intravascular catheter and denture biofilm formation [9], further emphasizing the concept that different TF networks and interactions are required for different types of *C. albicans* infections.

**Fig. 1.**
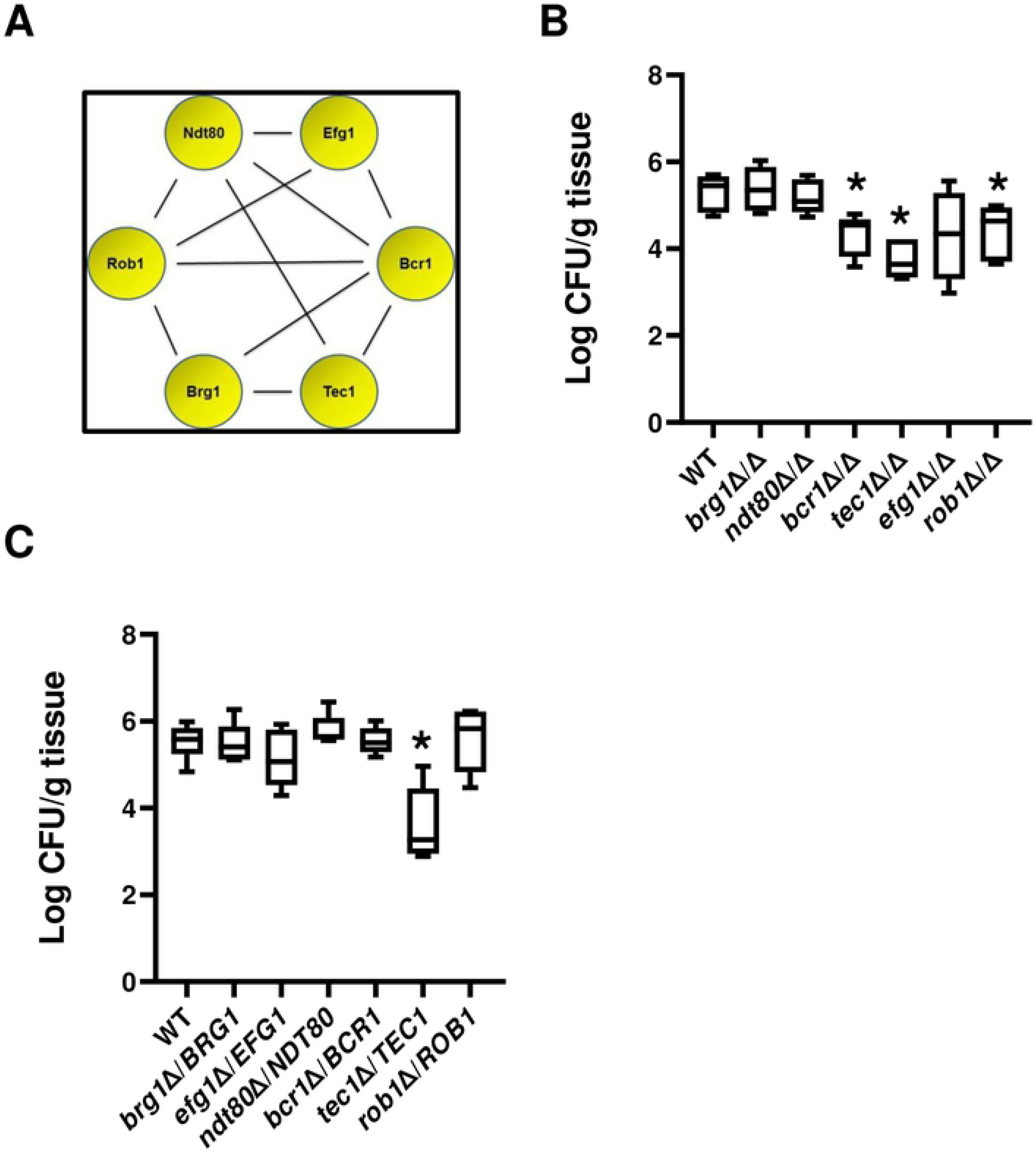
Effect of in vitro biofilm transcription factors in a mouse model of oropharyngeal candidiasis. A. The functional genetic interaction network of six transcription factors required for *C. albicans* biofilm formation in vitro [11]. B&C. The fungal burden five days post infection for animals infected with the indicated deletion mutants in the SN background. Median and standard deviation are shown for 4-5 animals per group. The log_10_-transformed fungal burden data for each experiment was analyzed by one-way ANOVA followed by post hoc Student’s t test to identify statistically significant differences between individual strains (*P* < 0.05). Strains that were statistically different than WT are indicated with an asterisk (*).

#### Complex haploinsufficiency-based analysis reveals critical interaction between Brg1 and Efg1 during murine OPC

Recently, we explored the genetic interactions between the 6 biofilm TFs in vitro using a set of all possible double heterozygous mutants derived from the core TFs [11]. To identify genetic interactions during murine OPC, we used this set to examine the ability of the double heterozygous mutants to infect the oral mucosae compared to the corresponding single heterozygote and WT strains. In contrast to the six negative genetic interactions that were identified within this network during in vitro biofilm formation [11], we found only one negative interaction which was between *brg1*Δ/*BRG1 efg1*Δ/*EFG1* (Fig. 2A, B). Interestingly, the reduction in fungal burden displayed by the *brg1*Δ/*BRG1 efg1*Δ/*EFG1* double heterozygote was similar to that shown by the *tec1*ΔΔ and *bcr1*ΔΔ mutants. Furthermore, mice infected with the *brg1*Δ/*BRG1 efg1*Δ/*EFG1* mutant had lower fungal burden than animals infected with the homozygous deletion mutants of either *EFG1* or *BRG1*, further establishing the interdependence of the function of these two TFs during murine OPC. In addition to the negative genetic interaction between *BRG1* and *EFG1*, we found that the haploinsufficiency of the *tec1*Δ/*TEC1* strain was suppressed in double heterozygotes containing *BRG1*, *EFG1*, *NDT80*, *BCR1*, and *ROB1* (Fig. 2A, C). The most common reason for these positive genetic interactions is that the two genes function in a linear pathway or trigger a compensatory response [13]. Based on these data, the functional genetic interaction network for these TFs during OPC can be formulated as shown in Fig. 2D.

**Fig. 2.**
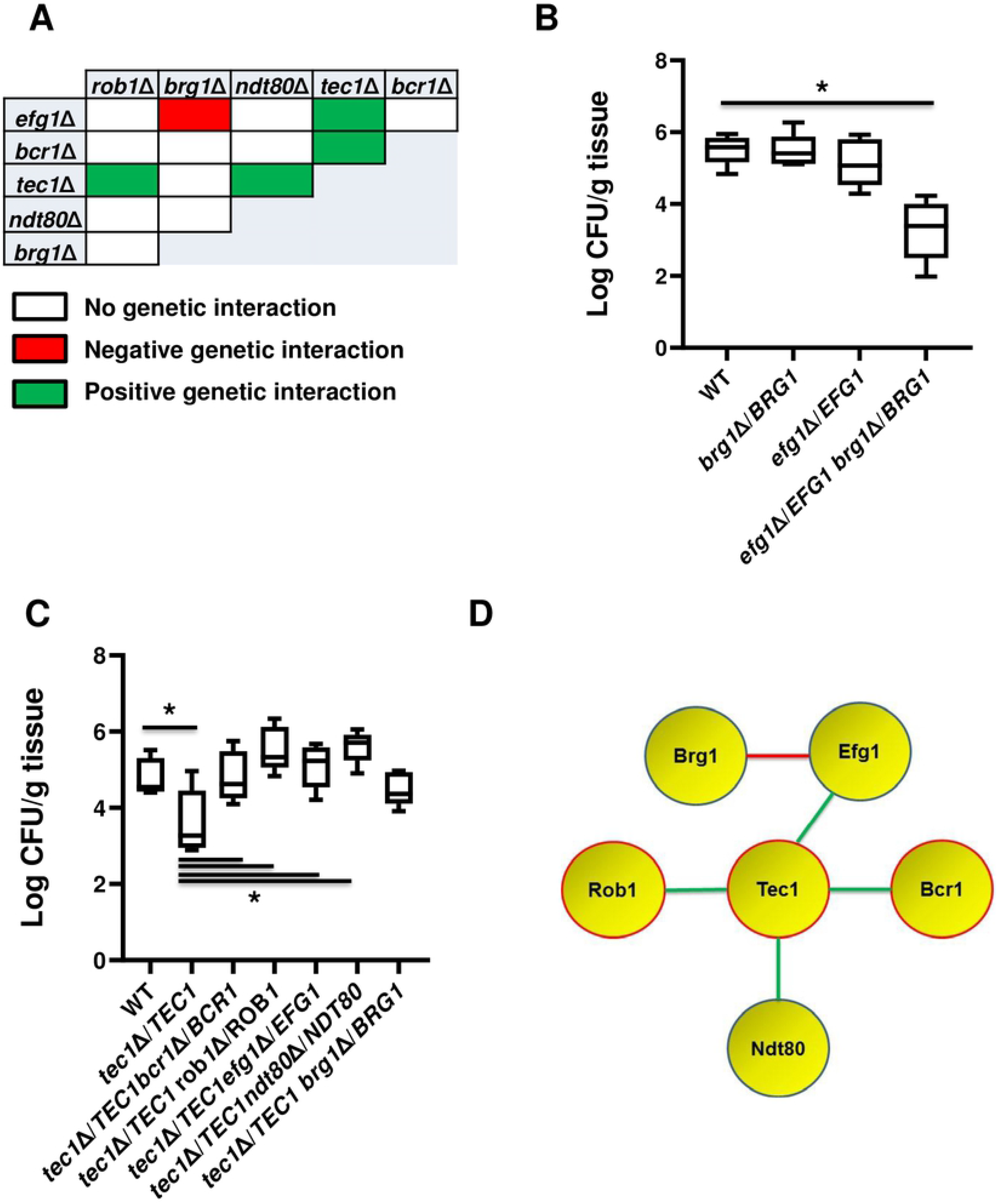
Genetic interaction analysis of transcription factors in a mouse model of oropharyngeal candidiasis identifies a functional network. A. The set of all possible double heterozygous transcription factor deletion mutants were screened for genetic interactions relative to the individual heterozygotes. The interaction map summarizes these interactions with negative interactions indicated in red; positive interactions indicated by green; and no interaction by white. B. The double heterozygous *efg1*Δ/*EFG1 brg1*Δ/*BRG1* mutant shows complex haploinsufficiency in the oropharyngeal candidiasis model. The log_10_-transformed fungal burden data for each experiment was analyzed by one-way ANOVA followed by post hoc Student’s t test to identify statistically significant differences between individual strains (*P* < 0.05). Strains that were statistically different than WT are indicated with an asterisk (*). C. *BCR1*, *ROB1*, *NDT80*, and *EFG1* show positive genetic interactions with *tec1*Δ/*TEC1*. D. The functional interaction network based on genetic interactions shown by the indicated transcription factors. Genes highlighted in red have reduced infectivity as homozygous deletion mutants. Red lines indicate a negative genetic interaction while green lines indicate a positive genetic interaction.

To further confirm the genetic interaction between *EFG1* and *BRG1*, we generated the double homozygote mutant, *brg1*ΔΔ *efg1*ΔΔ with the expectation that this strain would be even more severely affected than the *brg1*Δ/*BRG1 efg1*Δ/*EFG1* mutant. Consistent with that expectation, the tongues of mice infected with the double homozygous *brg1*ΔΔ *efg1*ΔΔ mutant were essentially sterile (Fig. 3A). Since the interaction between *EFG1* and *BRG1* was so significant in the murine model of OPC, we wondered if it was also manifest in a model of disseminated candidiasis. The *brg1*ΔΔ and *efg1*ΔΔ single mutants are both profoundly attenuated in the disseminated infection model so examination of the double homozygous strain would not be informative [14, 15]. We, therefore, compared the double heterozygote (*brg1*Δ/*BRG1 efg1*Δ/*EFG1*) to the single heterozygotes (*brg1*Δ/*BRG1* and *efg1*Δ/*EFG1*) and WT in the disseminated candidiasis model based on inoculation by tail-vein injection. As shown in Fig. 2B, the kidney fungal burden was slightly increased in animals infected with *brg1*Δ/*BRG1 efg1*Δ/*EFG1* double heterozygous mutant while the Kaplan-Meir curves for the four strains did not differ significantly (Fig. 3C). Thus, OPC infection is highly dependent on the genetic interaction between *EFG1* and *BRG1* while this interaction is much less important during disseminated candidiasis.

**Fig. 3.**
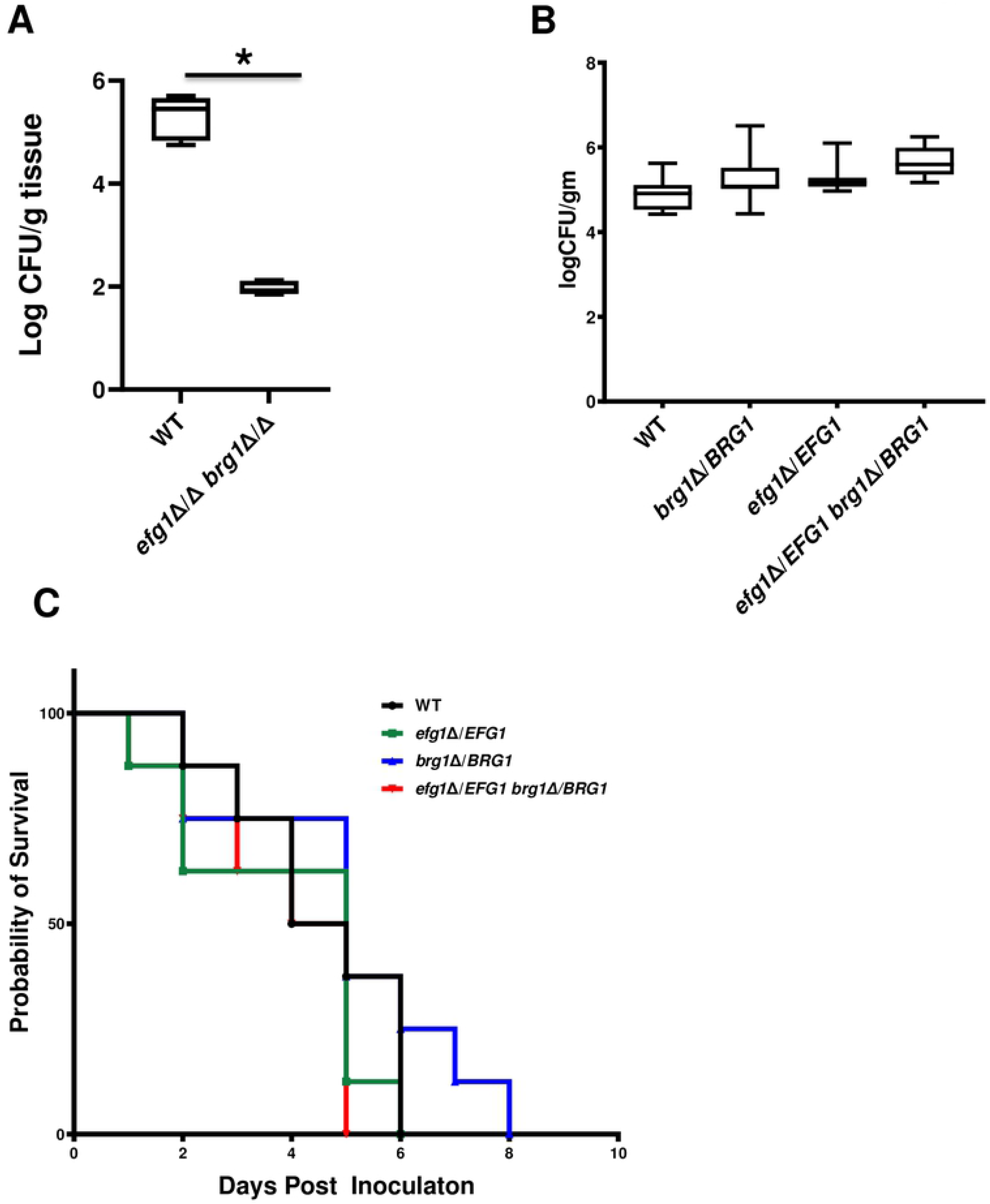
The genetic interaction of *EFG1* and *BRG1* is not observed in a model of disseminated candidiasis. A. The double homozygous *brg1*ΔΔ *efg1*ΔΔ shows a dramatic reduction in fungal burden relative to wild type in a mouse model of oropharyngeal candidiasis. The log_10_-transformed fungal burden data for each experiment was analyzed by one-way ANOVA followed by post hoc Student’s t test to identify statistically significant differences between individual strains (*P* < 0.05). Strains that were statistically different than WT are indicated with an asterisk (*). B. The double heterozygous *efg1*Δ/*EFG1 brg1*Δ/*BRG1* mutant does not have a reduced kidney fungal burden relative to WT or single heterozygous mutants in a mouse model of disseminated candidiasis. The log_10_-transformed fungal burden data for each experiment was analyzed by one-way ANOVA followed by post hoc Student’s t test to identify statistically significant differences between individual strains (*P* < 0.05). C. The double heterozygous *efg1*Δ/*EFG1 brg1*Δ/*BRG1* mutant is as virulent as WT and the single mutants in a mouse model of disseminated candidiasis based on Kaplan-Meier analysis of the disease progression curve shown.

#### Expression of *TEC1* from a heterologous promoter restores the infectivity of the *brg1*Δ/*BRG1 efg1*Δ/*EFG1* mutant but not *bcr1*ΔΔ

In addition to the single negative genetic interaction, we observed five positive or suppressive interactions involving the single haploinsufficient mutant, *tec1*Δ/*TEC1*. As noted above, positive interactions typically result when two genes function in a linear pathway [13]. During in vitro biofilm formation, the expression of *BCR1* is dependent on Tec1 while the expression of *TEC1* is partially dependent on Efg1 and Brg1 [9, 16]. Based on these molecular interactions, one model for the transcriptional circuit regulating OPC is shown in Fig. 4A. In this model circuit, *TEC1* is a key node and the model predicts that Brg1 and Efg1 play a critical role in the regulation of *TEC1* expression which, in turn, regulates *BCR1*. A second possible model is that Brg1 and Efg1 are redundant regulators of a set of genes that are independently regulated by Tec1 and Bcr1.

**Fig. 4.**
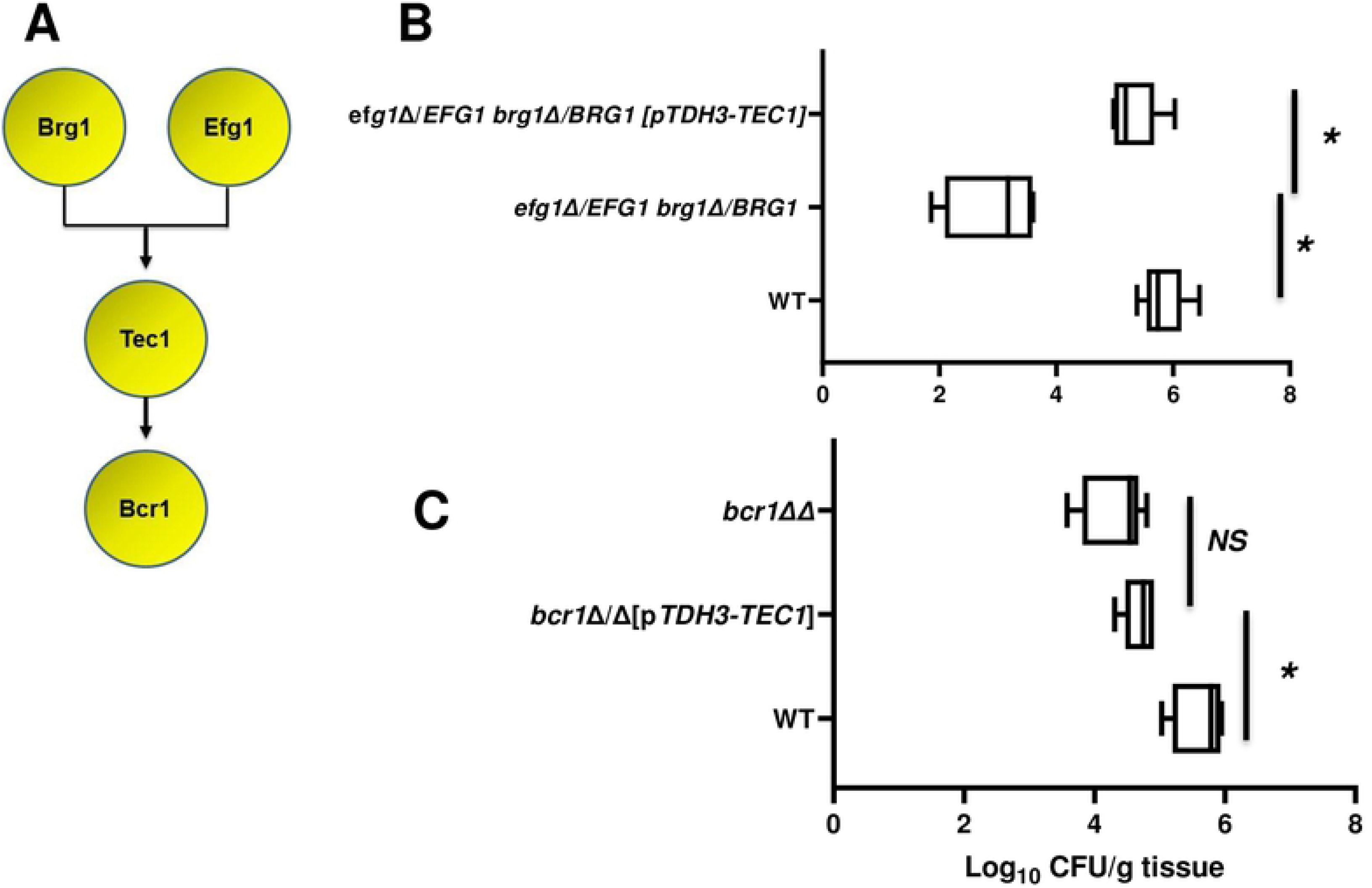
Epistasis analysis provides genetic support for a transcriptional circuit regulating infectivity in a mouse model of oropharyngeal candidiasis. A. Diagram of proposed regulatory circuit based on in vivo genetic interaction data and known binding interactions in vitro. B. Expression of *TEC1* from the *TDH3* promoter in the double heterozygous *efg1*Δ/*EFG1 brg1*Δ/*BRG1* mutant restores its infectivity in a mouse model of oropharyngeal candidiasis. The log_10_-transformed fungal burden data for each experiment was analyzed by one-way ANOVA followed by post hoc Student’s t test to identify statistically significant differences between individual strains (*P* < 0.05). C. Expression of *TEC1* from the *TDH3* promoter in the homozygous *bcr1*ΔΔ mutant does not restore its infectivity in a mouse model of oropharyngeal candidiasis. *NS* = not significant.

To test this model, we took a genetic epistasis approach. If *TEC1* expression was critically altered in the *brg1*Δ/*BRG1 efg1*Δ/*EFG1* double mutant during OPC infection, we reasoned that placing *TEC1* under the control of a promoter that is independent of Brg1 and Efg1 would increase the infectivity of the double heterozygous mutant [11, 12]. Consistent with that hypothesis, the *brg1*Δ/*BRG1 efg1*Δ/*EFG1* p*TDH3*-*TEC1* strain showed a 2.5 log_10_ increase in fungal burden relative to the parent double heterozygote (Fig. 4B). Previous in vitro data have shown that Tec1 regulates *BCR1* [16]. Our data indicate that both *TEC1* and *BCR1* are required for full OPC infectivity. The in vitro experiment does not, however, establish that Bcr1 is an effector of Tec1 and, thus, the question of whether the function of Tec1 is dependent on Bcr1 remained open. To test the latter hypothesis, we placed *TEC1* under the *TDH3* promoter in the *bcr1*ΔΔ background. As shown in Fig. 4C, p*TDH3*-driven *TEC1* did not alter the fungal burden defect of *bcr1*ΔΔ. These data are consistent with the model that Bcr1 functions as a Tec1 effector. However, it is important to note that it is possible that altered expression of *TEC1* could compensate for a decrease in Brg1-Efg1 input and, thus, the epistasis data do not rule out model number two.

#### Transcription factor mutants with decreased infectivity in murine OPC undergo filamentation in vivo

Five of the six TFs we examined have been shown to be required for filamentation in a wide range of in vitro inducing conditions [9,12]; *BCR1* is the exception in that is required for filamentation in some clinical isolates [17] and does interact with other TFs to modulate filamentation under some conditions [12]. Indeed, *EFG1* and *BRG1* have been described as master regulators of filamentation. Filamentation is required for biofilm formation and previous work has indicated that some strains with reduced filamentation also have reduced infectivity in murine OPC. Thus, one potential mechanism for the reduced virulence of the mutants identified above is that they have reduced filamentation following attachment to the epithelium.

We first examined the histology of tongues infected with homozygous deletion mutants (Fig. 5). Although Ndt80, Brg1, Rob1, and Efg1 are all known to be involved in the yeast-to-hyphae transition in vitro, none of the TF deletion mutants had significant defects in hyphal formation. When grown under hyphae-inducing conditions, *ndt80*ΔΔ mutants have a cell separation defect and form connected chains of pseudohyphae-like cells [18]. The tongue lesions formed by *ndt80*ΔΔ contain cells that have filamentous characteristics that are indistinguishable from those observed in lesions infected with WT (Fig. 5). Brg1 is a well-characterized regulator of hyphal morphogenesis in vitro and has been shown to play a key role in the maintenance phase of hyphal elongation [15, 19]. As with the *ndt80*ΔΔ mutants, the lesions on tongues infected with *brg1*ΔΔ mutants show robust filamentation. Likewise, lesions induced by the *rob1*ΔΔ mutants show a predominance of hyphae. *EFG1* is widely considered a “master-regulator” of filamentation in *C. albicans* [14, 20]. However, the fungal burden caused by *efg1*ΔΔ was not statistically different from WT; although we note that there was significant variability in the fungal burden in animals infected with *efg1*ΔΔ relative to other strains. The tongue lesions caused by infection with *efg1*ΔΔ mutants contained mainly filamentous fungal elements. *BCR1* does not affect filamentation in strains derived from SC5314 (e.g., the SN background, [17]) and, consistent with those observations, the *bcr1*ΔΔ mutant forms significant hyphae in the tissue.

**Fig. 5.**
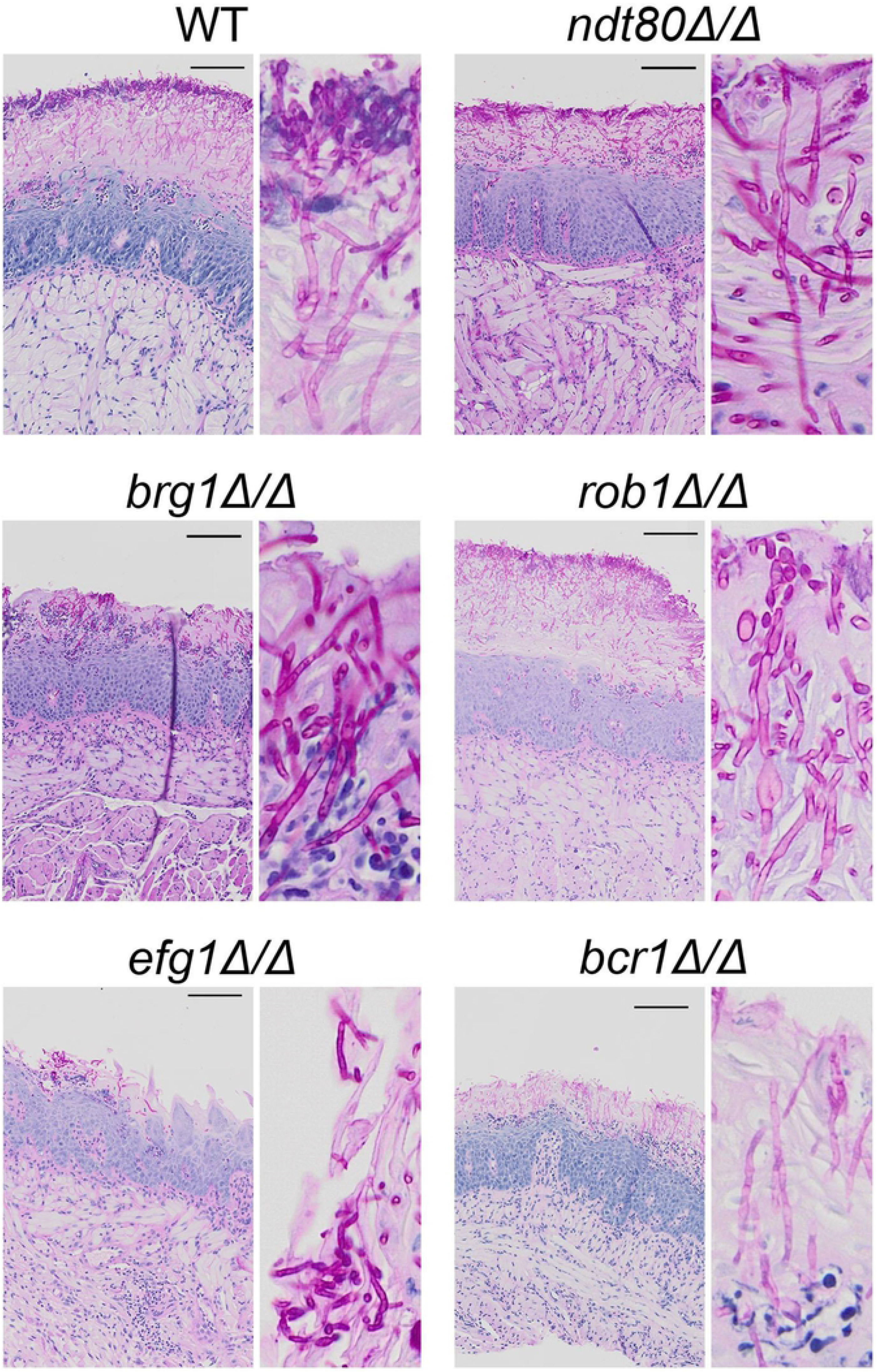
Homozygous transcription factor mutants with decreased infectivity undergo filamentation in tongue tissue. Histological sections of tongue tissue infected with the indicated strains were stained using the Periodic acid-Schiff (PAS) reagent. *C. albicans* organisms stain fuchsia in lesions. Insets are provided to show morphology.

Finally, the *brg1*Δ/*BRG1 efg1*Δ/*EFG1* double heterozygote had a lower fungal burden than mice infected with any of the single homozygote mutants. However, the lesions of the tongue from mice infected with this strain showed clear filamentation (Fig. 6). In vitro the *brg1*Δ/*BRG1 efg1*Δ/*EFG1* strain did not have significantly decreased proportions of filaments nor were the filaments shorter than WT or either single mutant length [12]. Although the double homozygous mutant *brg1*ΔΔ *efg1*ΔΔ has a very low burden in the tongue, we were able to identify scattered lesions in histological sections from tongues infected with this strain. It appears that this mutant also forms short filaments within the tissue of the tongues but it is difficult to make firm conclusions regarding this strain. Thus, somewhat surprisingly, *C. albicans* TF mutants with decreased infectivity in the mouse OPC model retain the ability to undergo the yeast to hyphae transition in vivo despite having significant filamentation defects in vitro.

**Fig. 6.**
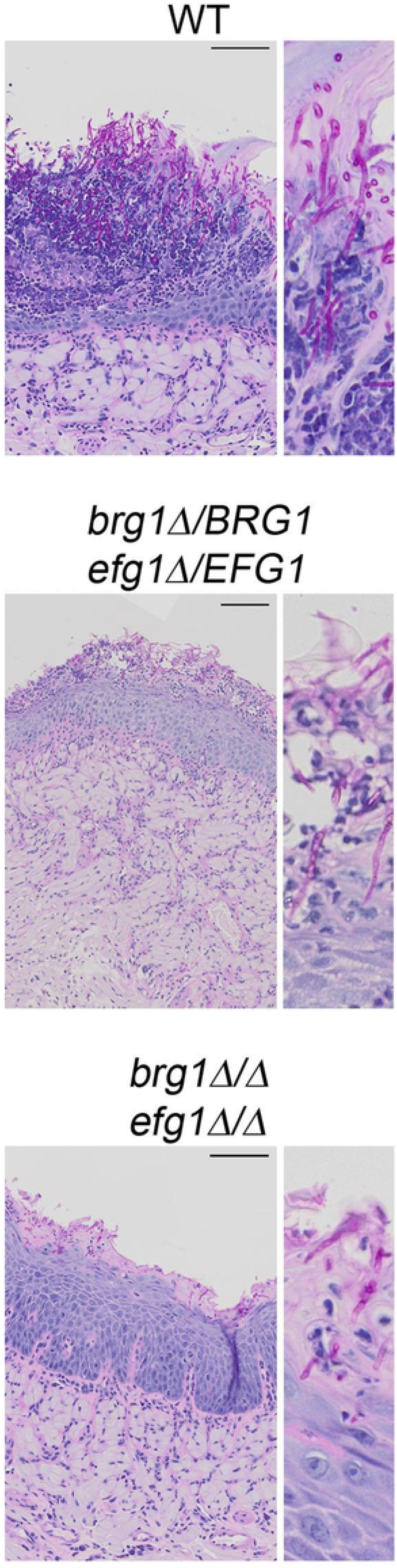
Double heterozygous and homozygous *EFG1*/*BRG1* mutants retain ability to filament in vivo. Histological sections of tongue tissue infected with the indicated strains were stained using the Periodic acid-Schiff (PAS) reagent. *C. albicans* organisms stain fuchsia in lesions. Insets are provided to show morphology

#### The Brg1-Efg1 interaction is required for oral epithelial cell invasion and is by-passed by constitutive expression of *TEC1*

In order to infect and damage oral tissue, *C. albicans* must adhere to and invade or be endocytosed by oral epithelial cells [21]. To determine which step of the process was dependent on the Brg1-Efg1 interaction, we used a well-established in vitro model based on the OKF6/TERT-2 oral epithelial cell line [22]. As shown in Fig. 7A, the *brg1*Δ/*BRG1 efg1*Δ/*EFG1* strain adheres to the epithelial cells as well as WT, but the double heterozygous mutant has a significant defect in endocytosis. Consistent with the data from the OPC murine infection model, constitutive expression of *TEC1* from the *TDH3* promoter in the *brg1*Δ/*BRG1 efg1*Δ/*EFG1* mutant restores endocytosis. These data strongly suggest that Brg1 and Efg1 regulate genes critically important for oral epithelial cell endocytosis/invasion and that these transcription factors function upstream of *TEC1* during OPC.

**Fig. 7.**
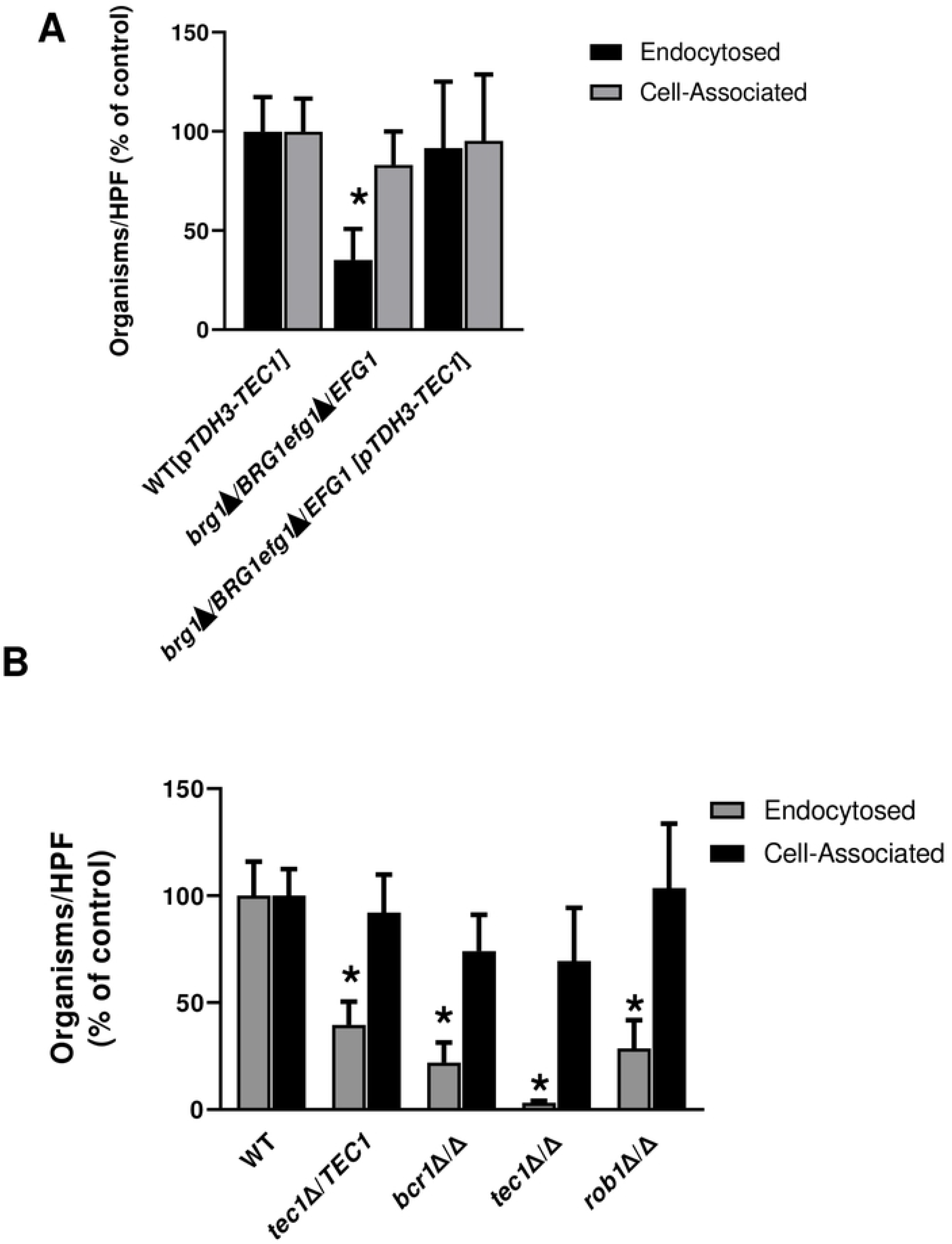
Transcription factor mutants with decreased infectivity in mouse model of oropharyngeal candidiasis have reduced induced-endocytosis of oral epithelial cells. A. The *efg1*Δ/*EFG1 brg1*Δ/*BRG1* double heterozygous mutant adheres to oral epithelial cells to an extent similar to WT but has reduces endocytosis. Bars indicate mean of two independent replicates and error bars are the standard deviation. Groups were compared by one-way ANOVA followed by post hoc Student’s t test to identify individual strains with statistically significant differences (*P* < 0.05: indicated by asterisk, *). B. Homozygous deletion mutants with reduced infectivity show deficits in oral epithelial cell endocytosis but not adherence.

Next, we examined the epithelial cell interactions of heterozygous and homozygous TF deletion mutants with decreased OPC fungal burden (Fig. 7B). Like the *brg1*Δ/*BRG1 efg1*Δ/*EFG1* double heterozygous mutant, the adherence of the haploinsufficient *tec1*Δ/*TEC1* cells to epithelial cells is similar to WT cells but its rate of endocytosis is decreased significantly; indeed, the *tec1*Δ/*TEC1* mutant is essentially quantitatively equivalent to the *brg1*Δ/*BRG1 efg1*Δ/*EFG1* strain. The homozygous *tec1*ΔΔ strain shows the same pattern as the heterozygous *tec1*Δ/*TEC1* mutant but the homozygous mutant is nearly completely unable to undergo endocytosis, clearly showing a gene dosage effect. This phenotype is also displayed by both the *bcr1*ΔΔ and *rob1*ΔΔ mutants. Interestingly, loss of Tec1 leads to a more severe phenotype than loss of the downstream TF, Bcr1, suggesting that Tec1 has Bcr1-dependent and -independent effects on epithelial cell endocytosis.

## Discussion

We have undertaken the first systematic genetic interaction analysis of *C. albicans* oral infection using a murine model of OPC. In doing so, we have identified two new transcriptional regulators of OPC, *TEC1* and *ROB1*. Using a complex haploinsufficiency, we have also elucidated a regulatory circuit (Fig. 4B) that is required for *C. albicans* to infect oral tissue and which appears to mediate the ability of *C. albicans* to undergo induced endocytosis when attached to epithelial cells. Finally, we find that TF deletion mutants such as *efg1*ΔΔ and *brg1*ΔΔ undergo filamentation during oral infection despite being largely unable to filament under most in vitro conditions, suggesting that the transcriptional regulation of this process is dependent upon the specific environmental niche.

We focused our screen of TF on a set previously shown to be highly interconnected during the process of in vitro and in vivo biofilm formation [9, 11]: *EFG1*, *TEC1*, *BCR1*, *BRG1*, *ROB1* and *NDT80*. The pathobiology of OPC appears to have features in common with in vitro biofilm formation [6]. Consistent with that model, previous studies have shown that, for example, the TF *BCR1* plays a critical role during in vitro biofilm formation, in vivo biofilm formation on oral dentures and intravascular catheters, and OPC in mouse models. Our results confirmed the role of *BCR1* in OPC [7, 8, 16] and found that *TEC1*, a regulator of *BCR1* expression in vitro [7, 9], is also required for full infectivity of *C. albicans* in this model. *ROB1* was the only other TF for which the homozygous deletion mutant showed reduced infectivity relative to WT reference strains. In vitro and during biofilm formation on in animate surfaces in vivo, all six of the TF mutants we examined show profound defects while during OPC only three of the mutants have significant phenotypes. This suggests that the functions of the interconnections between the TFs in this network are distinct during OPC.

One of the differences in our results compared to previous genetic analysis of OPC is that *efg1*ΔΔ mutants did not show a statistically significant effect [23]. Previous studies reported by one of us found that *efg1*ΔΔ mutants were less able to establish infection during OPC [23]. As shown in Fig. 1B, we observed significant variability in the fungal burden of mice infected with the *efg1*ΔΔ mutant. It is, therefore, possible that Efg1 affects OPC infectivity but it that would require larger sample sizes to confirm. It should also be mentioned that both the *C. albicans* and mouse strain backgrounds used in this work are different than that studied previously: SN vs. CAI for *C. albicans* and CD-1 vs Balb/c for mice [23]. As discussed below, we find that Efg1 does, in fact, play a role in OPC in conjunction with *BRG1*. Recent studies by Huang et al. have also shown that genetic circuits involving these very TFs can undergo significant diversification between different strain backgrounds in *C. albicans* [17]. It is, therefore, also possible that the slightly diminished role of *EFG1* during OPC indicated by our single gene deletion experiments results from subtle changes in network connections between different strain backgrounds as described by Huang et al. [17].

Previous reports by Nobile et al. [9] and Glazier et al. [11] have firmly established that this set of TFs regulate the expression and function of each of the other five TFs. During in vitro biofilm formation, this TF network is an example of a highly fragile genetic network because homozygous deletion mutants of any single TF are unable to form a biofilm [24]. Further supporting the fragility of the in vitro biofilm network, single heterozygous deletion mutants have reduced biofilm formation (i.e., haploinsufficiency) and double heterozygous mutants have phenotypes that are similar to homozygous deletion mutants during biofilm formation [11]. This is the opposite of a robust network where function is maintained in the face of multiple genetic disruptions [25]. An example of a robust phenotype is the fact that very few TF deletion mutants show decreased growth in standard yeast media (25).

Robust networks are combinations of “*or*” statements in which function is maintained **<u>if</u>**: this TF **<u>or</u>** that TF **<u>or</u>** this other TF are also functioning properly [24]. Fragile networks, on the other hand, are combinations of “*and*” statements in which function is maintained **<u>only if</u>**: this TF **<u>and</u>** that TF **<u>and</u>** this other TF are also functioning [24]. Our analysis of this same network of TFs during OPC infection indicates that it is more robust when compared to the fragile network characteristics displayed by the same mutants during in vitro biofilm formation. Supporting that conclusion are the following distinctions: **1)** 3 of 6 homozygous deletion mutants have reduced OPC infectivity while 6/6 have reduced in vitro biofilm formation; **2)** 1/6 heterozygous mutants showed haploinsufficiency during OPC but 5/6 were haploinsufficient during in vitro biofilm formation; and **3)** 1/15 combinations of double heterozygotes showed a negative genetic interaction in OPC infection while 6/15 displayed a negative genetic interaction during in vitro biofilm formation [11,24].

A trivial explanation for the apparent robustness of this network during OPC would be that some of the component TFs have no role in the establishment of infection. However, our data argue against this explanation. First, five of the TFs (*EFG1*, *BRG1*, *TEC1*, *BCR1*, and *ROB1*) are required for OPC infection either based on single or double mutant phenotypes. Second, the remaining TF, *NDT80*, shows a positive interaction with *TEC1*, suggesting it may function in a buffering role. Interestingly, the interaction between *TEC1* and *NDT80* during OPC is the opposite of the strong, negative complex haploinsufficient interaction observed during in vitro biofilm formation [11]. Overall, our genetic analysis of this network shows that its functional connections vary considerably between biofilm formation and OPC infectivity. The robustness of the latter may be due to the fact that *C. albicans’* natural niche is the oral cavity of mammals and, thus, it is more highly adapted to colonization of this niche than inanimate surfaces.

The single negative complex haploinsufficent interaction that we identified, *EFG1-BRG1*, combined with our observation that *TEC1* and *BCR1* are critical for full infectivity during OPC as well as previous observations in the literature outlined above allowed us to propose a potential genetic circuit (Fig. 4A). In vitro Tec1 regulates the expression of *BCR1* [16] and Brg1 and Efg1 are required for expression of *TEC1* [9]; additionally, all three TFs bind each other’s promoters during in vitro biofilm formation [9]. Hence, there is good literature precedent to support this model at the molecular level. Functionally, expression of *TEC1* from a promoter that is independent of *BRG1* and *EFG1* restored infectivity to the *brg1*Δ/*BRG1 efg1*Δ/*EFG1* strain while it is not sufficient to do so with the *bcr1*ΔΔ mutant (Fig. 4B/C). Thus, our genetic epistasis data strongly support the model and suggest that Bcr1 is likely to regulate key effector molecules involved in establishing infection. Previous studies support that model as well. Specifically, the overexpression of the Bcr1 target, *HWP1*, in a *bcr1*ΔΔ deletion mutant partially rescued OPC infection phenotypes [27]. However, it is important to note that, although Bcr1 is critical for regulating adhesins required for the establishment of biofilms on abiotic surfaces [7, 9], we found that *bcr1*ΔΔ mutants as well as the other mutants in the circuit were able to bind to an oral epithelial cell line similar to wild type (Fig. 5A/B). Thus, the decreased infectivity of the mutants in this circuit does not appear to be due to reduced ability of the fungus to bind the epithelial surface.

Instead, we found that the mutants showed a reduced ability to penetrate epithelial cells, a process mediated by induced endocytosis and/or direct penetration of the cells by hyphae [21]. The latter mechanism would appear to be an attractive explanation for some of our phenotypes since 5/6 components of the network are known to regulate hyphae formation in vitro. However, except in specific backgrounds, Bcr1 is dispensable for filamentation [17]. Furthermore, we were surprised to find that the homozygous and heterozygous deletion mutants with decreased OPC infectivity were able to undergo filamentation during infection based on histology of tongues infected with the mutants (Fig. 5). So, it seems unlikely that differences in hyphae formation and, thus, direct epithelial cell penetration can explain our results. Alternatively, it could be the case that the mutants fail to filament during the in vitro epithelial cell interaction studies and, thus, those studies could be confounded by differences in filamentation programs between in vivo and ex vivo conditions. We have, however, previously shown that the *brg1*Δ/*BRG1 efg1*Δ/*EFG1* filaments similarly to WT under tissue culture conditions [12] and, indeed, expression profiling of the mutant under those conditions focused on hyphae associated genes showed essentially no changes in expression in vitro (Do and Mitchell, unpublished results). Although a detailed understanding of the key outputs of this circuit during OPC awaits additional in vivo transcriptional profiling and characterization, it would appear that the circuit regulates genes that are important for mediating induced endocytosis.

Finally, our observations that in vitro regulators of hyphae formation are able to filament in the oral tissue represent an important finding. Most notably is the fact that mutants that lack two well-studied master regulators of filamentation, Efg1 and Brg1, form hyphae under these conditions. Although Efg1 is required for filamentation under a wide range of conditions [26], Riggle et al. have previously reported observing filamentous *efg1*ΔΔ mutants in oral infections of gnotobiotic pigs [28]. Thus, the oral cavity may provide filamentation stimuli that bypass the requirement for Efg1. Similarly, *BRG1* may also be bypassed under these conditions. It is possible that these two factors are redundant during filamentation but our limited histology of strains infected with the double mutant suggests filamentation is occurring to some extent. Although Tec1 is also required for filamentation under a variety of in vitro conditions [26, 29], it has been shown previously to form filaments in the tissue of kidneys mice in the setting of hematogenous candidiasis [29]. Thus, our observations are part of a growing body of evidence that the transcriptional regulators required for *C. albicans* filamentation are highly contingent on the specific strain and particular host or environmental niche [17, 30].

In summary, our systematic single gene and genetic interaction analysis of a set of TFs involved in OPC has revealed network and circuit level distinctions in how these TFs interact in vitro and in vivo. We have also identified a key transcriptional circuit that is required for infectivity and epithelial cell invasion. Future studies are needed to identify the key molecular outputs of this pathway and may identify novel mediators of *C. albicans* oral pathogenesis.

## Materials and Methods

### Ethics statement

All animal work was reviewed and approved by Institutional Animal Care and Use Committee (IACUC) of both the University of Iowa Carver College of Medicine and the Lundquist Institute at Harbor-UCLA Medical Center.

### Strains and media

The transcription factor (*EFG1*, *NDT80*, *BCR1*, *BRG1*, *TEC1*, and *ROB1*) heterozygous, homozygous, and double heterozygous deletion mutant strains are derived from the SN background and have been described previously. These libraries were screened in the OPC model as described below. As such, the strains were auxotrophic for arginine. TF mutants which showed decreased fungal burden in the OPC screening experiments were rendered prototrophic for arginine by transformation with StuI-digested plasmid, pEXpARG, which integrated the *ARG4* gene into the *RSP10* site [31] before confirmation of the phenotypes in repeat OPC experiments. The *TDH3* promoter was integrated into the 5’ region of *TEC1* in the *brg1*Δ/*BRG1 efg1*Δ/*EFG1* and *bcr1*ΔΔ by amplification of the promoter from plasmid pCJN542 [32] using a set of primers containing homology to the 5’ UTR and downstream of the T*EC1* ATG as previously described [11]. Strains were routinely streaked on yeast peptone 2% dextrose (YPD) plates from frozen stocks and incubated at 30°C. Prior to animal experiments or in vitro epithelial cell assays, the strains were grown overnight in liquid YPD with shaking at 30°C. The density of the culture was determined by hemocytometer and adjusted as indicated in the specific methods below.

### Mouse model of oropharyngeal candidiasis

The pathogenicity of the *C. albicans* strains were tested in the immunosuppressed mouse model of OPC as previously described with some modification [33]. Male ICR mice were injected subcutaneously with cortisone acetate (300 mg/kg of body weight) on days −1, 1, and 3 relative to infection. On day of infection, the animals were sedated with ketamine and xylazine, and a swab saturated with 106 *C. albicans* cells per ml was placed sublingually for 75 min. After 5 days of infection, the mice were sacrificed and the tongues were harvested. The harvested tongues were either homogenized and plated for quantitative fungal burden or sectioned and processed for histology. To limit the total number of mice, TF deletion sets were first screened with three animals per strain to identify mutants with statistically significant defects (*P* <0.05), or trends toward defects (*P* < 0.1) in the oral fungal burden. These mutants were then retested using five animals per group using strains that had been restored to arginine-prototrophy. The log_10_-transformed fungal burden data for each experiment was analyzed by one-way ANOVA followed by post hoc Student’s t test to identify statistically significant differences between individual strains (*P* < 0.05). Data reported in the figures are from the repeat experiments.

### Mouse model of disseminated candidiasis

Assessment of fungal burden and disease progression in a model of systemic candidiasis was performed using fully immunocompetent mice [34]. Female CD-1 outbred mice (6-8 weeks old; Envigo, Indianapolis, IN) were fed standard mouse chow or chow supplemented with 625 ppm doxycycline (Research Diets, Inc., New Brunswick, NJ) beginning three days prior to inoculation and throughout the experiment. Mice were then inoculated by lateral tail vein injection with 10^6^ colony forming units (CFU) of the indicated *Candida albicans* strains. For determination of kidney fungal burden, mice were euthanized 72 hours after inoculation. Kidneys were harvested, weighed, and homogenized in YPD. Ten-fold dilutions of the homogenates were plated in duplicate on YPD and incubated overnight at 30°C. The kidney fungal burden then was calculated as CFU per gram of kidney tissue homogenized. For evaluation of role of the *C. albicans* mutants in progression of disease, animals were inoculated as above and monitored daily for clinical changes. Any animal that demonstrated symptoms of severe disease (extreme fur ruffling, abnormal posture, difficulty with ambulation, failure to respond to surroundings) was euthanized immediately. Each experiment used 7-10 mice per group. Differences in the fungal burden were assessed using a one-way ANOVA followed by a post hoc Student’s t test using the log_10_ transformed data for fungal burden to identify statistically significant differences between strains (*P* <0.05). Disease progression was analyzed by Kaplan-Meier analysis and Log-Rank (Mantel-Cox test, *P* <0.05).

### In vitro assay of *C. albicans* adhesion and induced endocytosis with oral epithelial cells

To determine *C. albicans* adhesion to and invasion of oral epithelial cells we used our established protocol [22]. Briefly, OKF6/TERT-2 cells were grown to confluency on fibronectin-coated glass coverslips in 24-well tissue culture plates. The oral epithelial cells were infected with 2 × 105 yeast-phase *C. albicans* cells per well. After incubation for 2.5 hours, the cells were fixed with 3% paraformaldehyde, stained, and mounted inverted on microscope slides. The coverslips were viewed with an epifluorescence microscope, and the numbers of cell-associated and endocytosed organisms per high-power field was determined, counting at least 100 organisms per coverslip. Each experiment was performed at three times in triplicate. The means and standard deviations for each strain were determined and data were analyzed by one-way ANOVA followed by post hoc Student’s t test to identify individual strains with statistically significant differences (*P* < 0.05).

## Acknowledgements

The authors thank Alex Bassuk (University of Iowa) for access to imaging equipment used to generate histology figures. We also thank Martine Bassilana for providing plasmids.

## Author Contributions

Conceptualization: Damian J. Krysan, Scott G. Filler, Aaron P. Mitchell

Formal analysis: Norma V. Solis, Tomye L. Ollinger Melanie Wellington, Scott G. Filler, Aaron P. Mitchell, Damian J. Krysan

Investigation: Norma V. Solis, Rohan S. Wakade, Tomye L. Ollinger

Methodology: Rohan S. Wakade

Supervision: Damian J. Krysan, Scott G. Filler

Writing-original draft: Damian J. Krysan

Writing-review and editing: Damian J. Krysan, Scott G. Filler, Aaron P. Mitchell

